# Evolutionary conservation and divergence of human brain co-expression networks

**DOI:** 10.1101/2020.06.04.100776

**Authors:** WG Pembroke, CL Hartl, DH Geschwind

## Abstract

Mouse models have allowed for the direct interrogation of genetic effects on molecular, physiological and behavioral brain phenotypes. However, it is unknown to what extent neurological or psychiatric traits may be human or primate-specific and therefore, which components can be faithfully recapitulated in mouse models. We identify robust co-expression modules reflecting whole brain and regional patterns of gene expression and compare conservation of co-expression in 116 independent data sets derived from human, mouse and non-human primate representing more than 15,000 total samples. We observe greater co-expression changes occurring on the human lineage than mouse, and substantial regional variation that highlights cerebral cortex as the most diverged region. Cell type specific modules are the most divergent across the brain, compared with those that represent basic metabolic processes. Among these, glia are the most divergent, three times that of neurons. We show that regulatory sequence divergence explains a significant fraction of co-expression divergence. Similarly, protein coding sequence constraint parallels co-expression conservation, such that genes with loss of function intolerance are enriched in neuronal, rather than glial modules. We also identify dozens of human disease risk genes, such as COMT, PSEN-1, LRRK2, and SNCA, with highly divergent co-expression between mouse and primates or human. We show that 3D human brain organoids recapitulate *in vivo* co-expression modules representing several human cell types, which along with our analysis of human-mouse disease gene divergence, serve as a foundational resource to guide disease modeling and its interpretation.

## Introduction

The human brain is the culmination of millions of years of evolution (Sousa, Meyer, Santpere, Gulden, & Sestan, 2017), showcasing high level cognitive processes such as symbolic thought, self-reflection and the ability for long-term planning. Much of our understanding of human brain and disease is derived from studies in mice (Eppig et al., 2015). However, extrapolation from mouse models may be limited because humans last shared a common ancestor with mice 90 million years ago (mya) (Hedges, Dudley, & Kumar, 2006; Mouse Genome Sequencing et al., 2002). Humans exhibit a five-fold relative expansion in cortical volume, changes in cellular composition, and cell-type differences, such as a substantial increase in size and complexity of human astrocytes (Herculano-Houzel, Mota, & Lent, 2006; Hodge et al., 2019; Oberheim et al., 2009). Furthermore, many distinctions, such as the molecular reorganization of specific cell types and neural circuits have arisen over recent human evolution (Sousa, Meyer, et al., 2017). A number of neurological disorders are associated with the dysfunction of biological processes that occur in brain regions or cell types that possess features specific to human biology (Miller, Horvath, & Geschwind, 2010; Skene & Grant, 2016; Wang et al., 2016), posing the question as to which components underlying human neuropsychiatric diseases can be faithfully recapitulated in model organisms, or for that matter, cell-based model systems (Muchnik, Lorente-Galdos, Santpere, & Sestan, 2019).

Human evolution is hypothesized to be driven primarily by changes in gene regulation rather than protein sequence divergence (Chimpanzee & Analysis, 2005; King & Wilson, 1975; Konopka et al., 2012), highlighting the transcriptome as an appropriate nexus for investigating evolution, as demonstrated by recent studies assessing gene expression in brains across multiple primate species (Sousa, Zhu, et al., 2017; Zhu et al., 2018). While many gene expression changes may be a result of neutral drift over the course of evolution (Khaitovich et al., 2004), gene co-expression networks provide a functional framework for assessing whether changes in expression are indeed neutral, or have a functional impact (Oldham, Horvath, & Geschwind, 2006). Gene co-expression networks built on data from human brain tissues have been utilized to assess which aspects of human brain function are preserved and diverged across species (Konopka et al., 2012; Miller et al., 2010; Oldham et al., 2006). These studies were either conducted using small sample sizes from a single, or a few brain regions, or constructed “brain-wide” co-expression networks, which limits the identification of more subtle “intra-region” transcriptomic patterns.

Due to natural variation in cell-type proportion across tissue samples, co-expression analysis using homogenous tissue with large sample sizes allows us to generate cell-type information without the need to physically isolate cells, so called “in silico dissection” (Kelley, Nakao-Inoue, Molofsky, & Oldham, 2018; Oldham et al., 2008). Single-cell sequencing of both human and mouse cortex has allowed identification of species differences at a cell-type resolution (Hodge et al., 2019). However, this analysis was limited to a single brain region and focuses on differential gene expression within each cell class. Because co-expression reflects functional mechanisms such as co-regulation, changes in network position reflect changes in function (Oldham et al., 2006; Parikshak, Gandal, & Geschwind, 2015). Therefore, investigating co-expression networks from multiple brain regions across different phylogenetic groups may allow us to assess the functional divergence of biological processes and cell types for many brain regions.

To identify robust evolutionary divergent brain regions and biological processes we constructed co-expression networks for 12 brain regions in human based on the GTEx resource (Consortium et al., 2017; Hartl et al., 2020) and 7 brain regions in mouse from 30 studies (Table S1, S2). We assessed network divergence for each brain region in human and mouse by performing module preservation in the corresponding brain region for human, non-human primate (NHP) and mouse. Our analysis was in line with previous findings, with glial co-expression modules on average three times as divergent as neuronal modules (Kelley et al., 2018; Miller et al., 2010). By exploring many major regions across the brain, we were able to identify regional variation and compare preservation across regions. We observed that cerebral cortical brain regions display the greatest relative divergence in human, an analysis enabled by the extensive regional sampling within the GTEx dataset (Consortium et al., 2017). We identify 5,473 genes that display significant divergence of co-expression from human to mouse in at least one brain region, including many that have been related to human neurological and psychiatric disease. By examining the relationship of co-expression divergence with measures of divergence at the DNA sequence level, including regulatory sequence changes and protein coding sequence constraint (pLI; ref), we identify genetic mechanisms underlying divergence and show that the extent of transcriptomic divergence reflects other known measures of selection. Since co-expression modules are often related to cell-type (Kelley et al., 2018; Oldham et al., 2008), these divergent genes can begin to explain the nature of cell-type divergence between human and mouse. We show that gene divergence in co-expression is associated with changes in mean expression between species. Through integration with single-cell sequencing data, we show that these species differences in gene expression are largely due to cellular differences in gene expression rather than cell type proportion differences in bulk tissue. Of these diverged genes, 91 (2%) show evidence of genetic association to at least one neurological disorder, such as the autism (ASD) risk genes SCN2A and SHANK3. Furthermore, a substantial proportion (18%) of genes that display up- or down-regulation in post-mortem brain of patients with neurological disorders display significant divergence of co-expression from human to mouse, potentially limiting their use as disease biomarkers in mouse.

## Results

### Assessing the evolutionary divergence of human and mouse brain networks

We generated regional co-expression networks in a discovery dataset for both human and mouse to identify modules of highly co-expressed groups of genes (Fig 1A; Methods). Networks were created using a robust bootstrapping approach to ensure modules were not driven by outlier samples (Langfelder, Luo, Oldham, & Horvath, 2011). To validate whether these modules represent generalizable co-expression relationships, we performed module preservation analysis against multiple independent expression datasets derived from human (3-20 studies, depending on region; 7,287 total samples), non-human primate (NHP) (5-15 studies; 2,933 total samples) and mouse (6-28 studies; 6,667 total samples; Methods) to assess co-expression conservation (Fig 1B; Table S1,2). We determined quantitative module-level co-expression differences between species (Methods, Fig 1B) to assess the divergence of different brain regions, cell types and the genes which underlie this divergence (Fig 1B, C). We reasoned that modules highly conserved in multiple independent data sets from one species (e.g. human), but not in the independent data sets from another, were divergent between the species (Methods).

**Figure 1.**
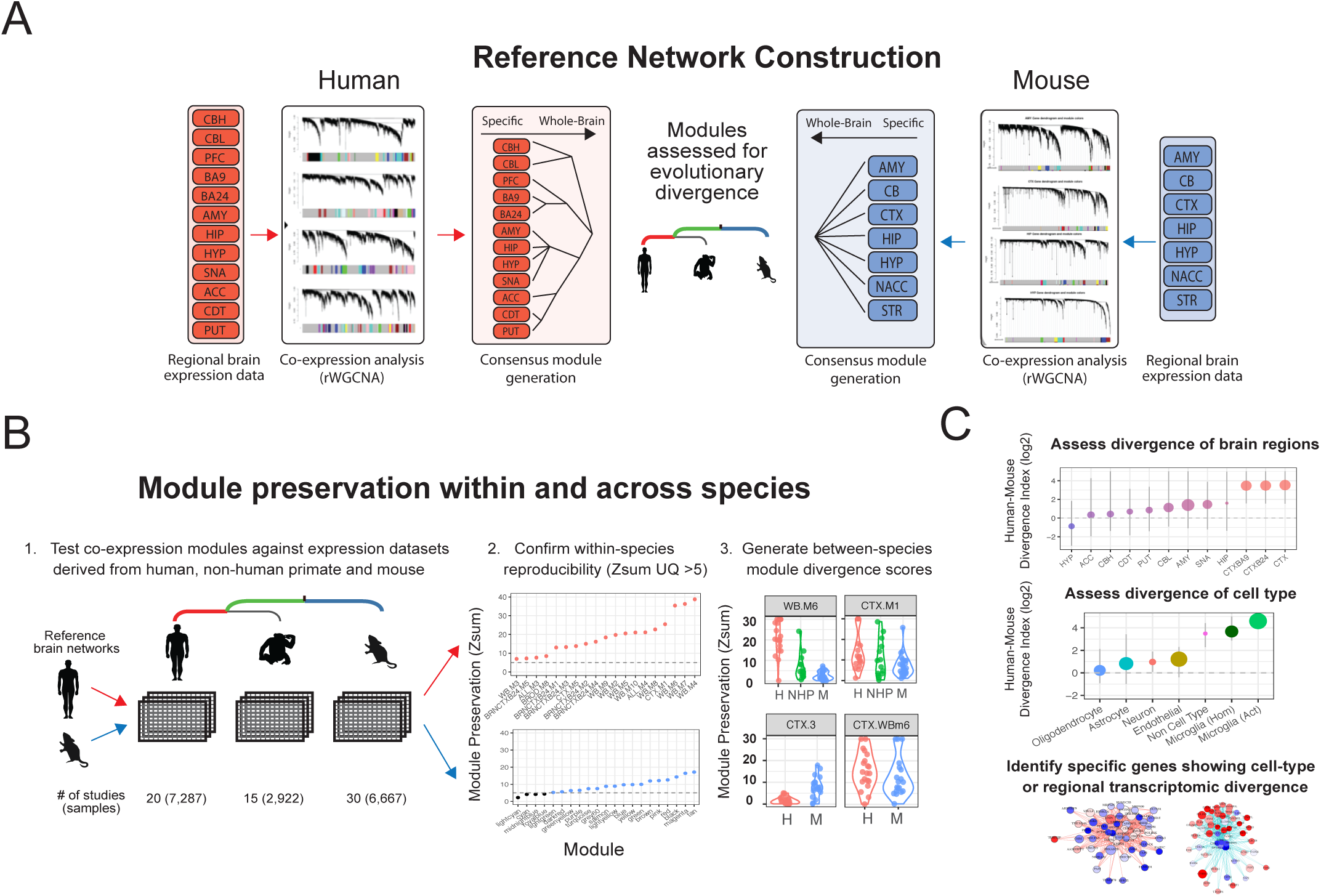

### Co-expression analysis in human and mouse reveals asymmetric transcriptome divergence across brain regions

To capture divergence in co-expression relationships between the species, we derived divergence scores for human and mouse modules. Module divergence scores were calculated by comparing the module preservation (Zsum) scores from test datasets of the same species (eg. human) to the Zsum scores of test datasets derived from the opposing species (eg. mouse) (Fig 2A). Human modules, on average, displayed over twice the divergence than modules defined in mouse (Methods; OR=2.5, p < 1e^-6^), suggesting an “asymmetric” transcriptomic divergence, with more changes occurring on the human lineage (Fig 2B). The preservation of most mouse modules in human suggests that core transcriptional programs in mouse are shared with human. However, since many human modules show divergence from mouse, there may be biological processes in human with additional levels of transcriptomic complexity that are not captured in mouse.

**Figure 2.**
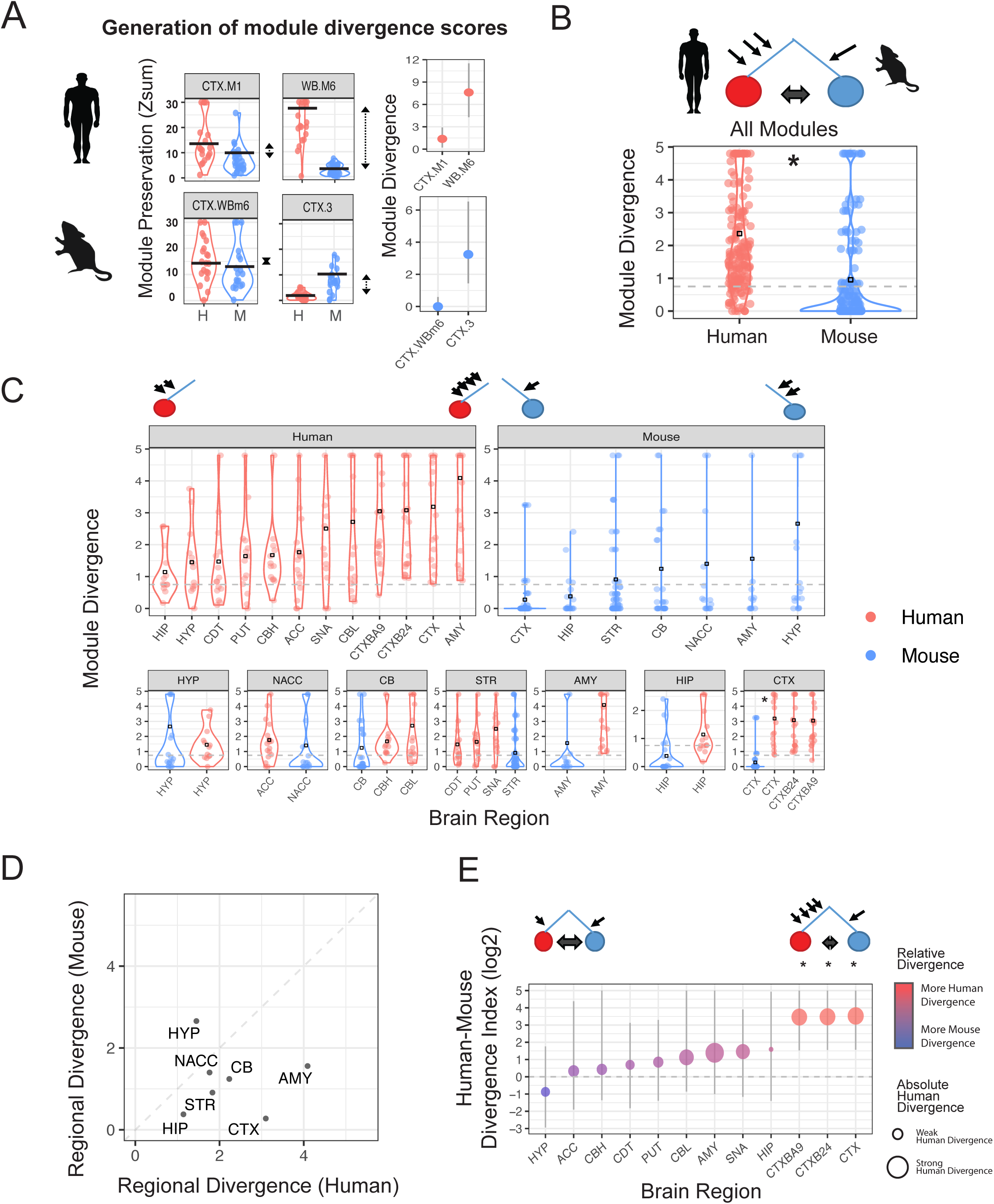

We explored this notion of asymmetric divergence further by stratifying module divergence according to brain region, allowing us to assess regional divergence in co-expression. We observe that the cerebral cortical regions show the greatest asymmetric divergence; human derived, but not mouse-derived cortical modules were more diverged in the opposing species (p < 1e^-3^; Fig 2C,E). On the other hand, the cerebellum shows similarly minimal divergence for both human and mouse, suggesting this region has not strongly diverged for either species (Fig 2C-E). The amygdala and hypothalamus both show substantial divergence (> 1.4 average module divergence) in *both* human and mouse, meaning that there are several modules that are divergent in each region in each species. While these regions possess a small degree of asymmetric divergence, this does not reach significance (Fig 2C-E). We confirmed that this regional level trend of module divergence was not confounded by sample size, or the number of studies contributing to that network (Fig S1A). Another potential confounding factor would be the variation in the proportion of different cell-type modules across the brain regions. We tested this by normalizing regional divergence, so that the proportion of different cell-type modules were equal across regions, and found that regional divergence was not driven by the regional proportion of cell-type modules (Fig S1B).

### Glial cell types display the greatest transcriptomic divergence in human

To further understand which biological processes were preserved or diverged across species, we tested modules for their enrichment in cell-type marker gene lists and gene ontology terms (Methods; Fig S1C-D,S2A-B). The cell-type classification of each human module was highly associated with human-mouse module divergence (p < 1e^-10^; Kruskal Wallis), with glial-classified modules significantly more divergent than neuronal (OR > 3.1; p < 1e^-6^; pairwise t-test; Fig 3A-B). Human microglial modules show the largest divergence (mean = 4.8), followed by astrocytes (4.3), oligodendrocytes (2.9), and neurons (1.4). In contrast, modules with no evidence of cell-type enrichment were generally well preserved in mouse (mean = 1.1). This category of highly conserved, non-cell-type specific modules included those enriched for GO terms relating to ribosomal-related processes, RNA-binding and energy production (Fig S2C).

**Figure 3.**
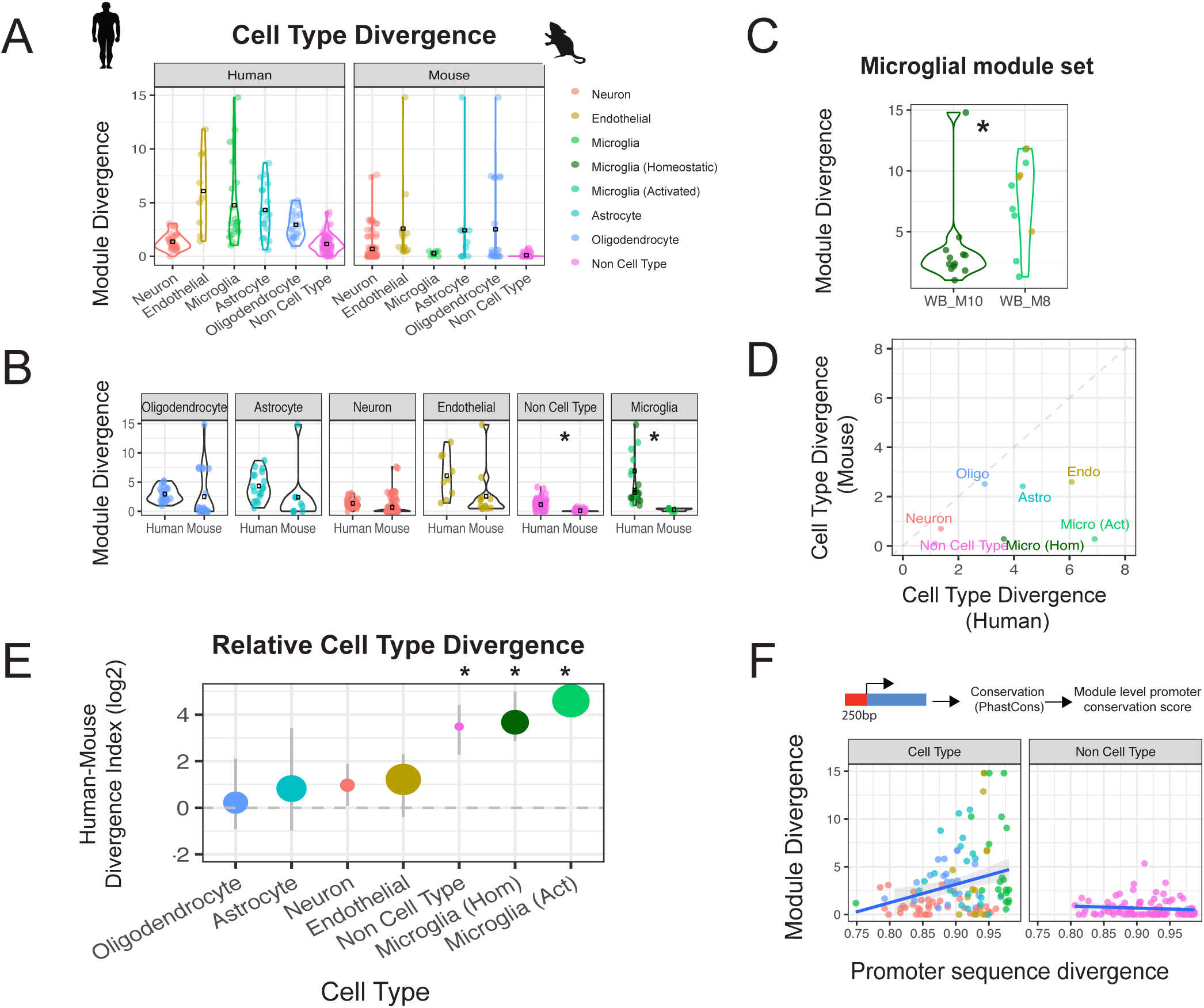

To confirm that the difference in divergence between different cell types is not related to network construction, we performed preservation analysis using collated cell-type marker gene lists derived from co-expression and single-cell experiments (Methods; Fig S2A) and found a strong correlation with our cell-type module divergence scores (Fig S2D; Pearson’s cor = 0.69 (co-expression) and 0.88 (single-cell)). To assess whether species cell type differences may arise from differential aging trajectories (Galatro et al., 2017; Loerch et al., 2008), we tested module preservation in young adult (human 13-40 years; mouse 2-14 months) and middle age to aging adult (human >40 years; mouse >14 months) samples (Fig S2E; Methods). Preservation scores in human are similar between young and aging adult brain indicating that co-expression modules are not driven by age-dependent processes (Fig S2E-F). Preservation was also largely similar between aging states in mouse (Fig S2E-F). Notable exceptions were human microglial modules WB.M8 and WB.M10, which displayed stronger preservation in aging mice than young mice (OR>1.8; pval<1e ^-10^; Fig S2F), and are described in more detail below.

Human microglial marker genes largely fall into two module subclasses, WB.M8 and WB.M10. These modules are generated utilizing all 12 brain regions and thus are referred to as whole brain (WB) modules (Fig 3C). To provide more refined annotations of these modules, we assessed module-enrichment using microglial marker gene lists derived from immunopanned microglia, and brain tissue from both homeostatic and pathological conditions ((Holtman et al., 2015; Mancarci et al., 2017; Zhang et al., 2014; Zhang et al., 2016); Methods). Based on this, M10 represents a more canonical microglial signature defined by strong enrichment in canonical homeostatic microglial signatures (Fig S2G), whereas M8 represents a more activated microglial state, as defined by enrichment of endothelial genes and microglial genes upregulated in pathological conditions, such as Nfkbia and Cxcl1 ((Holtman et al., 2015); Fig S2G). Differences between these two states explain much of the variation observed for microglial module divergence, with the activated microglial modules (WB.M8) displaying stronger divergence between species (OR=4; p<0.01; Fig 3C). We also observe that there is a range of preservation of these microglia modules in mice, with relatively higher preservation in older versus young adult mice (Fig S2E-F). Nevertheless, the preservation in older mice is still much less than preservation in independent human datasets (human to mouse preservation OR=4.0, M8; OR=2.0, M10), indicating that a large proportion of microglial differences are consistent across all ages.

In addition to showing the greatest absolute divergence from human to mouse, glial cell classes also displayed strong asymmetric divergence. Human-derived microglial modules, on average, showed the most such divergence: over ten-fold greater divergence to mouse than mouse-derived microglial modules to human (Fig 3B,D-E). Stratification of modules by cell-type in each brain region revealed that in cerebral cortex, all cell types displayed significant asymmetric divergence, i.e. human-derived modules from all cell types were significantly more diverged to mouse than their corresponding mouse-derived cell-type modules were to human (Fig S2H). In contrast, most cell types across other brain regions did not show significant asymmetric divergence (Fig S2H).

### Transcriptomic divergence is highly associated with regulatory sequence divergence for cell-type modules

The divergence of co-expression from human to mouse could be driven by many factors, genetic or environmental. Matching environmental variables between the two species is not experimentally feasible, so to understand potential mechanisms underlying divergence, we focused on the extent to which genetic variation could explain the differences in co-expression observed between species. We reasoned that we could assess the relationship of co-expression divergence to the divergence in relevant DNA sequences directly by comparing core regulatory sequence divergence to module divergence. We correlated the average PhastCons score for a number of definitions of the most proximal promotor region (250b, 2kb upstream of all the genes in a module) with the module divergence score. Indeed, we observe a significant correlation (p < 0.01, Pearson’s cor = 0.27) between transcriptome and sequence divergence (Fig 3G) with neuronal modules displaying greater evolutionary constraint than glial modules, in line with recent findings (Hu, Li, & Wang, 2020). This transcriptome-sequence divergence correlation is maintained across a range of gene boundary parameters as a basis for sequence comparisons (Fig S3A-B), demonstrating that a cell-type specific selection pressure is not being driven by our parameter selection. Finally, we find that this correlation of regulatory sequence-transcriptome divergence is restricted to cell-type specific modules, rather than the more general metabolic, or housekeeping annotated modules, as the correlation between sequence and expression divergence is lost when assessing modules without cell-type enrichment (Fig 3G). For example, despite strong promoter divergence of ribosomal related modules, these processes appear to be highly resistant to transcriptomic perturbation (Fig S3C).

### Co-expression preservation is highly associated coding sequence constraint in cell-type modules

We next asked, given the positive relationship between module divergence and regulatory sequence divergence, whether other measures of selective pressure, such as protein coding constraint in humans was related to module divergence. Genes vary in their tolerance to loss of function (LoF) mutations on one allele (Lek et al., 2016) and can be classified depending on their frequency of spontaneously arising (*de novo*) mutations in the human population (Samocha et al., 2014). For genes where LoF mutations are well tolerated, the observed number of *de novo* mutations equals the expected number of mutations under the null model. Conversely, haploinsufficient genes, where half the total level of a gene product is insufficient for organismal survival, the observed *de novo* mutation frequency is defined to be <10% than what is expected under the null model (Lek et al., 2016). The pLI (probability of LoF Intolerance) score is the probability that a given gene is haploinsufficient and therefore intolerant to LoF variation and under purifying selection. We asked whether haploinsufficiency was evenly distributed across modules or related to the degree of co-expression divergence, finding a significant negative relationship between the degree of transcriptomic divergence and module enrichment for LoF intolerant genes (pLI ≥ 0.9; Pearson’s cor = -0.2; p < 0.01; Fig S3D) (Samocha et al., 2014). Modules with limited transcriptomic divergence, such as neurons, manifested the greatest enrichment in genes intolerant to LoF mutations. As functionality of these genes is necessary for organismal survival, we would expect these genes to be under tighter transcriptional regulation. Conversely, glial modules are under-represented for LoF intolerant genes, which is consistent with the relaxed constraint of these genes at the sequence and transcriptional level. This analysis, coupled with the regulatory sequence analysis above, further supports the functional significance of module divergence by demonstrating its relationship to independent measures of selection and constraint at the DNA sequence level.

### Identification of modules displaying accelerated divergence after the last common ancestor (LCA) with non-human primates

Understanding transcriptome divergence between human and mouse is essential due to the ubiquitous nature of mice in biomedical research (Eppig et al., 2015). However, it is also of interest to know if certain diseases are caused by disruption of biological processes changing most during human evolution, or whether they can be modelled in non-human primates (NHP), or human stem cell systems (Konopka et al., 2012; Qiu & Li, 2017). Comparison of the Zsum scores for human, NHP and mouse allowed the Human-Mouse divergence score to be partitioned into a 1) ‘human-specific’ component, where transcriptional changes occurred on the human lineage after divergence with the last common ancestor (LCA) of NHP, and a 2) ‘primate-specific’ component, where changes occurred before divergence with the LCA of NHP (Fig 4A-C).

**Figure 4.**
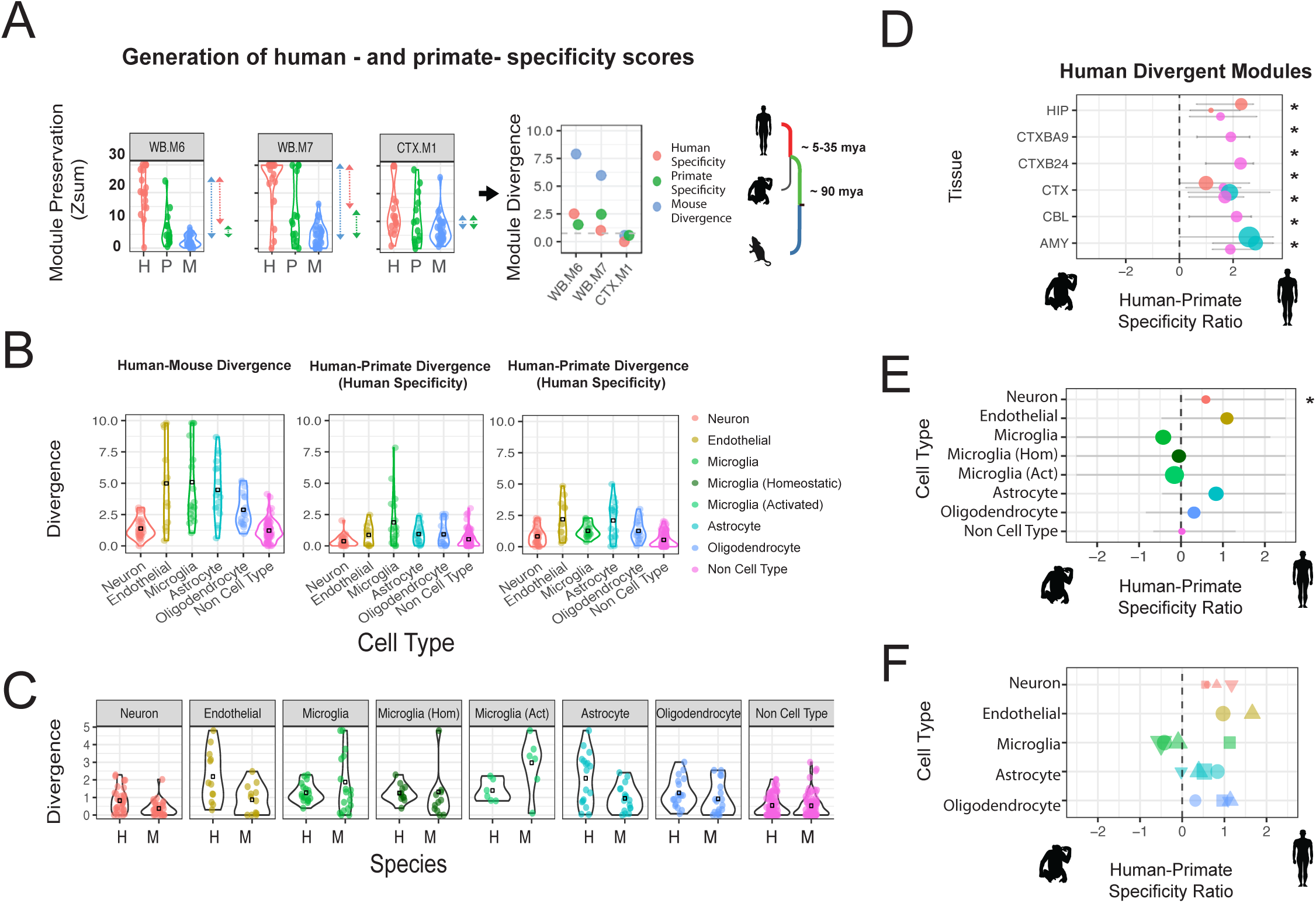

To quantify whether module divergence was greater before or after the LCA with primates, we compared the human- and primate-specificity scores to each other, producing a Human-Primate ‘Specificity Ratio’ (Fig 4D; Methods). If the preservation scores in NHP are closer to the preservation scores in mouse relative to human, it suggests more changes have occurred *after* the LCA of NHP, on the human-specific lineage. Conversely, if the preservation scores in NHP are closer to the preservation scores in human relative to mouse, it suggests more changes have occurred *before* the LCA of NHP, on the primate lineage. We were able to identify 13 co-expression modules where the NHP preservation scores were significantly closer to mouse relative to human and therefore displayed stronger divergence on the human-specific rather than primate lineage (Fig 4D). We were unable to identify any modules where the NHP module preservation scores were significantly closer to human than mouse, which would have been indicative of increased divergence on the primate lineage before the NHP - human divergence.

By grouping together modules from the same cell-type, and assessing preservation in NHP relative to human and mouse, we can determine the extent to which certain cell types have diverged before or after the LCA of NHP. Cell-type modules, apart from the microglial modules, showed slightly greater transcriptional changes on the ‘human-specific’ portion of the lineage tree, with the neuronal cell-class displaying significance (pval < 0.05; Fig 4B-E). This human-favored specificity ratio trend was also observed when preservation analysis was performed using cell-type marker gene lists derived from three different approaches: co-expression, cell sorting to obtain purified cell populations, and single-cell based methods (Fig 4F).

### Specific genes showing cell-type or regional transcriptomic divergence

To identify gene drivers of module divergence, we calculated the correlation of each gene to its module eigengene (kME) in the discovery dataset and all test datasets in human, NHP and mouse. The difference in mean kME values between human and mouse was used to calculate a kME divergence score, which can be utilized to highlight genes whose expression is highly divergent between human and mouse (Methods). To highlight genes underlying cell-type divergence, we calculated the kME divergence for each gene in the consensus ‘Whole-Brain’ modules across all brain regions. ‘Whole-Brain’ consensus modules were created from a co-expression network generated by creating a consensus of all regional co-expression networks (Methods) and therefore represent shared features of co-expression across the brain. Overall, we identified 2,217 genes that displayed consensus divergence across all 12 brain regions in human and 528 genes that displayed consensus divergence across all 7 brain regions in mouse (Table S3-S4). For human, we identified hundreds of genes whose co-expression is highly divergent for microglia (295; WB.M8/WB.M10), astrocytes (281; WB.M6), oligodendrocytes (272; WB.M7), neurons (469; WB.M4), and endothelia (88; WB.M11) (all listed in Table S3). These include 1,109 cell type associated genes which have not been previously determined to be divergent between mouse and human (Kelley et al., 2018; Miller et al., 2010), such as *TMIGD3, C3AR1* (microglia; WB.M10), *MID1, PDLIM3* (astrocyte; WB.M6), *CD22, FAM124A* (oligodendrocyte; WB.M7), *JAKMIP1 and NDUFA5* (neuron; WB.M4; Table S3). To illustrate human-mouse cell-type divergence, network plots display the top 40 most divergent genes for each consensus cell-type module (Fig 5A). As expected, highly divergent modules possessed a greater proportion of divergent genes (Pearson’s cor =0.48, p<1e^-11^; Fig S4A). Supporting our methodology, we find that previously identified human- and mouse-specific cell-type markers display significantly stronger kME divergence than background (Fig S4B; Human; OR=4.1; p<1e^-13^; Mouse; OR= 14.5; p<0.01 (Kelley et al., 2018); Human; OR=2.2; p<1e^-4^ (Miller et al., 2010)).

**Figure 5.**
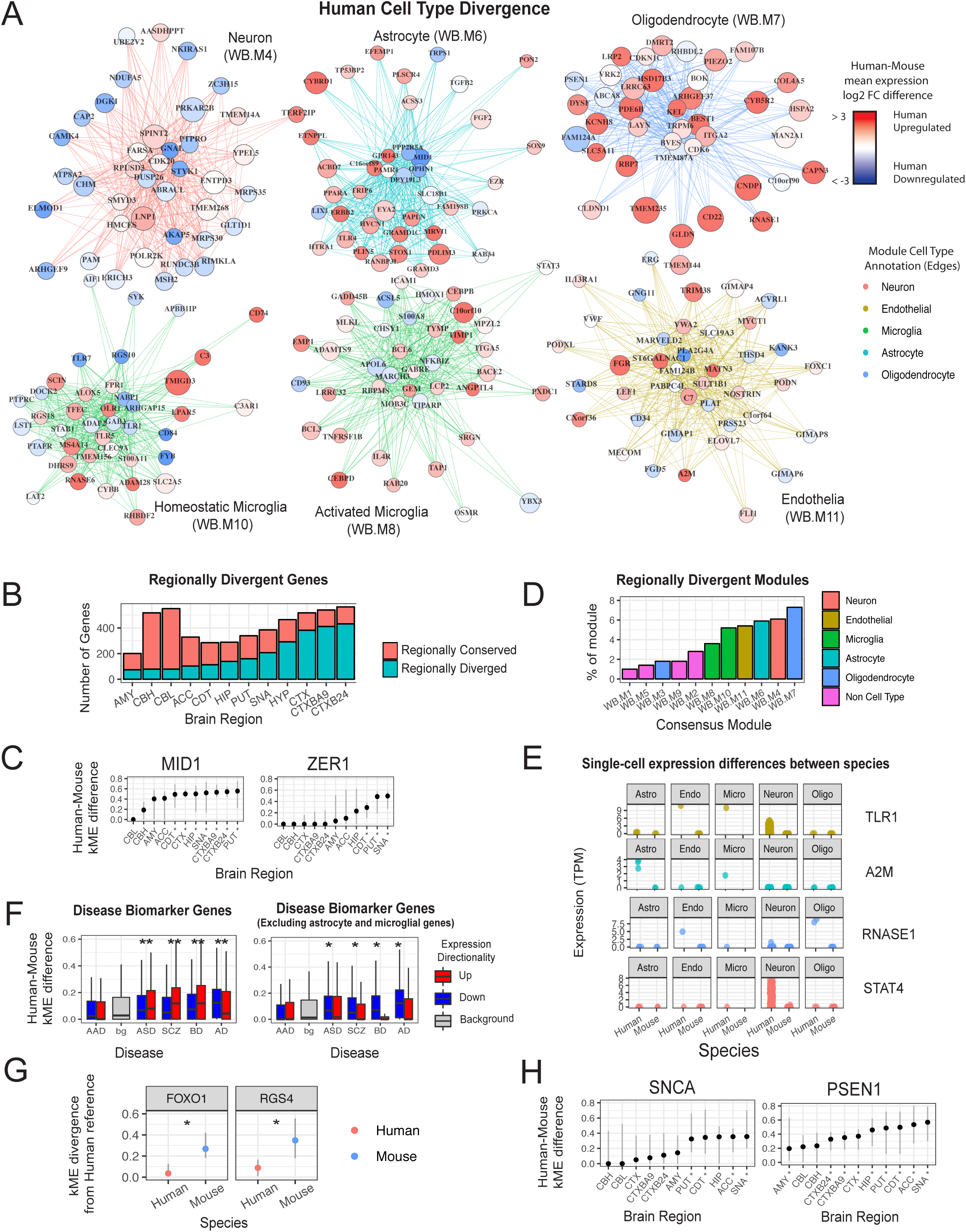

To highlight human-specific gene associations for each cell type, we compared gene-gene correlations in all human and mouse datasets. We selected the top 100 gene pairs from each ‘Whole Brain’ consensus module (Methods) that showed the strongest correlation differences between human and mouse (Table S5) (Kelley et al., 2018; Miller et al., 2010). As expected these genes show strong kME divergence (mean kME div = 0.17; OR=2.3; p<1e^-16^) with the strongest effect seen in glial modules (Fig S4C). For example, in oligodendrocyte module WB.M7, the recently identified Alzheimer’s risk gene HSPA2 (Petyuk et al., 2018) shows 7 times stronger human specific association with a number of oligodendrocyte markers in human than in mouse (Fig S4D), in line with a previous study (Miller et al., 2010). For microglial module WB.M10, in addition to C3, which has previously been identified as a human specific microglial marker (Kelley et al., 2018), we show that the complement cascade genes C1QA-C and C3AR1 form 44 of the top 100 highly divergent gene pairs. This strongly implicates a human-specific role for complement mediated synaptic pruning in microglia. Glutamate transporters SLC1A3 and SLC1A2 (EAAT1/EAAT2) show strong divergence with other members of the astrocytic WB.M6 module suggesting *human* astrocytes have an increased capability to provide glutamate to adjacent neurons.

We also identify 1,135 region-specific divergent genes, where kME divergence is restricted to particular brain regions (Table S6). The cerebral cortex displayed the greatest region-specific divergence with over 300 genes significantly more diverged than at least one other brain region (Fig 5B). For example, MID1, a member of the astrocytic WB.M6 module shows significantly stronger kME divergence in cerebral cortical regions than cerebellum (Fig 5C). The cerebellum displayed strong region-specific conservation (Fig 5B). This strong regional conservation is likely a combination of the general conservation of cerebellum from human to mouse (Fig 2E) and the large transcriptional differences of the cerebellum to the rest of the brain (Hartl et al., 2020; Kang et al., 2011; Zhu et al., 2018). Furthermore, genes comprising the neuronal (WB.M4) and oligodendrocyte (WB.M7) modules show the greatest region-specific divergence, with 6.1% and 7.3% of genes within each module respectively, displaying significantly greater divergence in a particular brain region (Fig 5D). This suggests that neurons and oligodendrocytes manifest the most divergence between the species on a region-specific basis.

We sought to test the hypothesis that human divergent genes would have greater expression in humans, relative to mouse. We assessed whether kME diverged genes were associated with changes in mean expression between human and mouse, finding that genes with greater expression in human (>4 FC) show greater divergence (OR=1.4; pval<1e ^-8^; Fig S4E). For example, highly diverged genes in the oligodendrocyte (WB.M7) and astrocyte (WB.M6) module appear to be strongly upregulated in human compared with mouse (Fig 5A, S4E), suggesting that up-regulation of gene expression is one mechanism driving higher co-expression.

To assess whether this change in expression levels between species is due to a change in gene regulation, or simply a change in cell proportion, we compared human-mouse expression differences from our cortical bulk expression data to single-cell expression data derived from human and mouse cortex (Hodge et al., 2019). Genes with species-differences in bulk-data that maintain differences in single-cell expression data are likely to be driven by species differences in gene regulation, whereas genes with species-differences only in bulk-data may be driven by differences in cell-type proportion. For cell types captured in both human and mouse single-cell datasets, there is a strong positive correlation between bulk- and single-cell data for the species differences in gene expression (Fig S4E). This indicates that cell-type proportion is not a major driver of these expression differences, but rather expression differences are largely due to cellular gene expression differences. We illustrate the single-cell expression data for four genes with greater expression in human for both bulk and single-cell expression data, indicating intracellular upregulation in human for a particular cell-type (Fig 5E). Divergent neuronal genes (WB.M4) generally displayed downregulation in bulk expression data, which was mirrored in single-cell data, indicative of an intracellular downregulation of neuronal genes in human (Fig S4F-G). This matches known age-dependent repression of neuronal genes on the primate lineage (Loerch et al., 2008).

Genes with divergent co-expression from human to mouse were more likely to show upregulation in both human single-cell and human bulk expression datasets. Nevertheless, a large portion (73%) of significantly divergent genes showed similar or lower expression in human (Fig S4E), indicating that changes in baseline expression level is not the sole mechanism. In addition, many genes that displayed strong expression differences between human and mouse in both single-cell- and bulk-datasets, did not show a strong divergence of co-expression (Fig S4E). This trend was especially true for downregulated genes - 15% of human-upregulated genes (> 2 FC) in both single-cell and bulk data did not display significant divergence in co-expression, whereas 66% of human-downregulated genes (< 0.5 FC) did not show significant co-expression divergence. This suggests that species differences in mean expression can partially predict species differences in co-expression, but there is a notable proportion of genes where this trend does not apply, further indicating that differential expression is only one mechanism driving differential co-expression (Amar, Safer, & Shamir, 2013; Farahbod & Pavlidis, 2019).

### Gene drivers of transcriptomic divergence in disease

Mouse models are widely used to understand the molecular and cellular mechanisms underlying human disease (Eppig et al., 2015). Therefore, we reasoned that understanding which disease-associated genes were conserved for co-expression between human and mouse would help inform disease modeling. Genes up- and down-regulated in human cortex from schizophrenia (SZ), bipolar disorder (BD), autism (ASD) and Alzheimer’s (AD), but not alcoholism (AAD) are significantly enriched for diverged genes (p < 0.01; Fig 5F) (Gandal et al., 2018). For example, FOXO1 and RGS4 are up- and down-regulated respectively in the cerebral cortex of patients with SZ, BD, ASD or AD (Mirnics, Middleton, Stanwood, Lewis, & Levitt, 2001), but their co-expression is highly diverged in mouse (Fig 5G). The divergent co-expression of these genes between species suggests they contribute to different biological processes in mice and humans (Kelley et al., 2018; Miller et al., 2010), potentially limiting their study or use as disease biomarkers in mouse. We next assessed whether the divergence of these disease-related expression changes was related to specific cell types or biological process. Indeed, genes upregulated in ASD, SCZ, BD and AD largely represent transcriptomically divergent glial and immunological signatures, as enrichment of kME divergent genes in upregulated genes for these disorders is attenuated when omitting genes from astrocyte or microglial annotated modules (Fig 5F).

Many transcriptional changes in post mortem tissue may be a consequence of disease rather than causal, prompting us to focus on genes with evidence of causality in human disease (Methods). We first examined enrichment of high confidence ASD risk genes (n=136), defined by harboring high risk likely protein-disrupting mutations (Ruzzo et al., 2019; Sanders et al., 2015; Satterstrom et al., 2020) of which SHANK3, SCN2A, and 56 other genes displayed significant kME divergence in at least one brain region (Table S7). In total, 70 ASD risk genes were divergent when including genes with relatively high levels of statistical support (within SFARI categories 1 or 2; n=170 (Abrahams et al., 2013). Of the genes with an association to a neurodegenerative disease (n=84), 40 displayed significant kME divergence from mouse (Table S7). For example, alpha-synuclein (SNCA), a Parkinson’s (PD) risk gene, shows co-expression divergence between human and mouse primarily in the substantia nigra and basal ganglia, whereas presenilin-1 (PSEN-1), an AD risk gene, which had been shown to be divergent in cortex (Miller et al., 2010) displayed significant divergence across 10 brain regions (Fig 5H; Table S7). We supplement these gene lists by assessing the divergence of genes within relevant KEGG disease pathways or the DisGeNet database and provide these as a resource to guide disease modeling in mouse (Methods; Table S8).

Leveraging NHP expression data, we were able to identify 1670 genes with human specific co-expression changes (strong kME divergence in human with respect to both NHP and mouse) such as the astrocyte gene, ACBD7 and the neuronal, PD risk gene SYT4 (Table S7,8). One thousand six hundred and sixty genes displayed primate-specific co-expression changes (strong kME divergence from human and NHP to mouse; Table S9) of which 23 had high-confidence disease association. This provides guidance as to genes that would be more aptly modeled in NHPs than mouse and suggests that modeling these disease genes in mouse should be approached with caution when attempting to relate mechanisms identified in mouse to primates or humans. These genes include the ASD risk gene SCN2A, the PD risk gene COMT and the AD risk gene PSEN-1 (Table S7), which has previously been shown to be divergent in its co-expression between human and mouse cerebral cortex (Miller et al., 2010).

### Identifying modules strongly preserved in human cortical spheroid models

Recent advances in *in vitro* modeling offer the potential to model human brain development and function in a dish (Kanton et al., 2019; A. M. Pasca et al., 2015; Sloan et al., 2017; Yoon et al., 2019). To better understand the fidelity of *in vitro* modeling for specific cell types, we assessed cerebral cortex module divergence for canonical cell-type modules in human cortical organoids derived from 8 separate studies (Birey et al., 2017; Camp et al., 2015; Kanton et al., 2019; Pollen et al., 2019; Pollen et al., 2015; Sloan et al., 2017; Trujillo et al., 2019; Velasco et al., 2019; Yoon et al., 2019). We then compared module divergence scores between mouse, NHP and organoids to assess which species and systems can recapitulate human neurobiology. Species “divergence scores” largely represent transcriptomic differences shaped by evolution, whereas organoid “divergence scores” reflect differences between *in vivo* and *in vitro* systems. Nevertheless, these scores can still be used to assess the applicability of using particular species or systems to model human *in vivo* signatures.

In general, astrocytic, activated glial and most neuronal modules were well captured in cortical organoids. Conversely, homeostatic microglial and oligodendrocyte modules were not (Fig 6A-C), which is expected given their absence from most organoid cultures (Kanton et al., 2019; Yoon et al., 2019). As previously shown, all cell types were generally well preserved in NHP datasets (Fig 6A-C). Mouse data recapitulated modules with neuronal or no cell-type annotation, but displayed strong divergence for glial cell types (Fig 6A-C).

**Figure 6.**
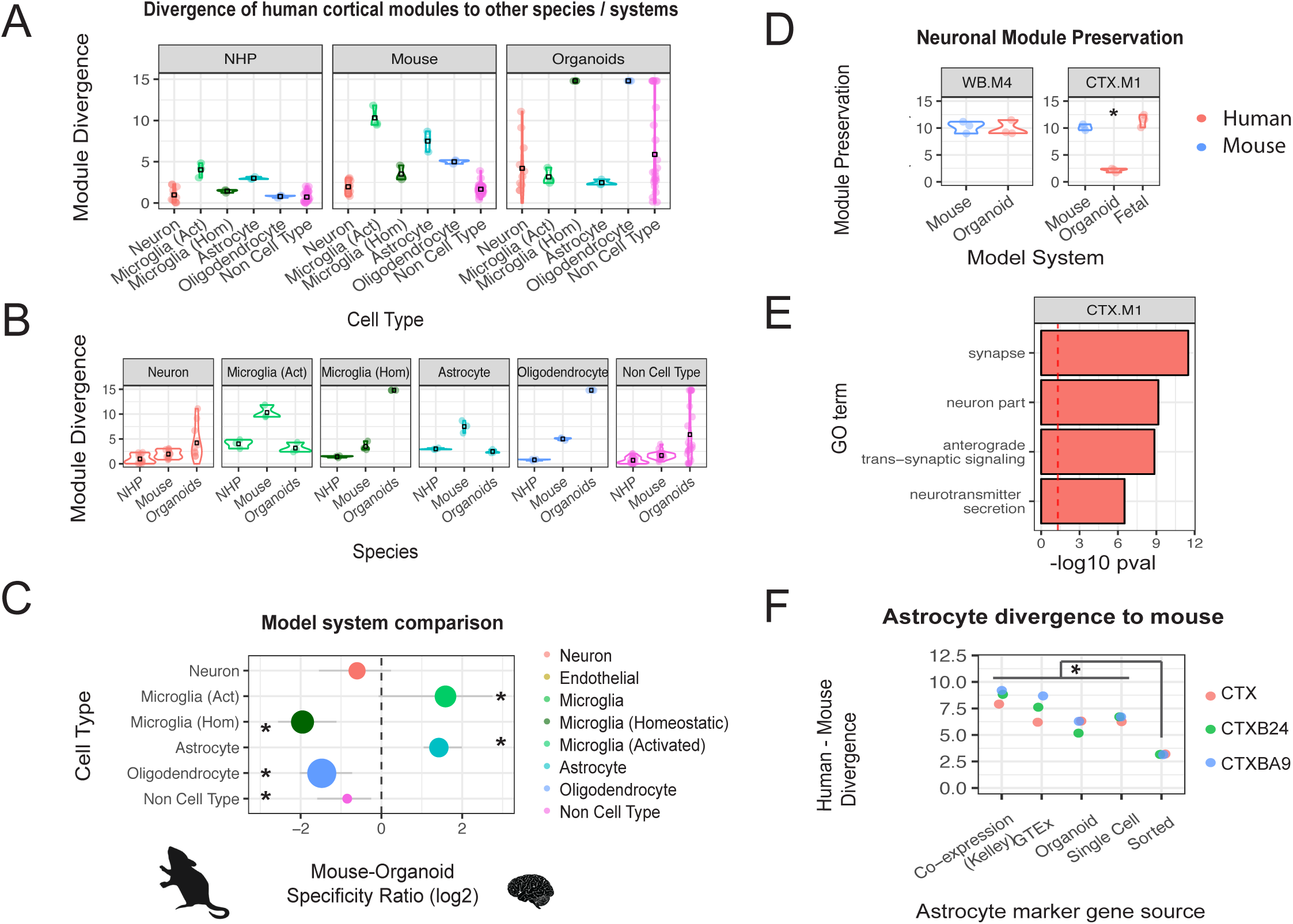

To summarize whether *in vivo* mouse or *in vitro* human approaches more appropriately model *in vivo* adult human cell types, we calculated the log fold change between the mouse and organoid module divergence scores. Modules relating to astrocytes displayed significantly greater preservation in human organoid cultures (Fig 6B,C), whereas modules relating to homeostatic microglia and oligodendrocytes displayed significantly greater preservation in mouse (Fig 6B,C), which is not surprising since most of these published IPSC models do not include these cell types (Marton et al., 2019). Neuronal modules, which were generally well preserved in mouse, displayed different degrees of co-expression preservation in organoids. A canonical cross neuronal-subtype module, WB.M4, shows strong preservation in organoids (Fig 6D). In contrast, the cortical module CTX.M1, which displays enrichment for excitatory pyramidal genes and ontology terms relating to synaptic transmission and vesicle transport (Fig 6E), displayed weak preservation in organoids despite showing preservation in mid-gestational developing human brain samples (Fig 6D; (Kang et al., 2011; Walker et al., 2019)). Therefore, although basic neuronal cell types are present *in vitro*, mature synaptic transmission pathways are not, most likely due to their immature state, as previously shown in 2D cultures (S. P. Pasca, 2018; Stein et al., 2014). It is clear that model systems need to be improved, as none of the mouse or organoid data analyzed here indicated that they faithfully recapitulate human oligodendrocyte and microglia transcriptional signatures (Fig 6B), consistent with recent studies focused on comparing organoid development to *in vivo* human datasets (Kanton et al., 2019; Pollen et al., 2019).

To confirm that organoids possess human-specific expression signatures not preserved in mouse, we assessed co-expression preservation in human and mouse using human astrocyte markers derived from organoid (Sloan et al., 2017), and from co-expression (this study & Kelley at al., 2018), single-cell (Lake et al., 2018) and immunopanned human brain (Zhang et al., 2016). Human organoid-derived astrocytes were highly divergent to mouse, similar to that of human astrocyte markers derived from co-expression or single-cell sequencing (Fig 6F). Interestingly, astrocyte markers derived from immunopanned human brain were largely lacking in human-specific co-expression patterns and displayed divergence scores significantly lower than the other astrocyte marker gene lists (OR=0.54, p<0.05). This suggests that the human-specific component of astrocytes may be highly dependent on its physiological environment (Khakh & Deneen, 2019), as taking astrocytes out of their 3D environment reduces their divergence to mouse. As human-derived astrocytes appear to lose human-specific expression signatures after immunopanning and culturing, previously observed species-specific differences using this methodology may have underestimated true *in vivo* divergence (Cahoy et al., 2008; Zhang et al., 2014; Zhang et al., 2016).

## Discussion

Several previous studies have used gene networks to identify genes whose co-expression has human specific features (Konopka et al., 2012; Miller et al., 2010; Oldham et al., 2006), usually focusing on a single or small number of brain regions due to data availability. Here, we perform the first comprehensive multi-region, multi-species comparison of the evolutionary divergence of co-expression networks generated from human brain derived from over 100 individuals (Hartl et al., 2020). Conservation of co-expression was tested in over 15,000 samples from 116 studies derived from human, NHP and mouse (Table S2). Comparison of transcriptomic conservation on a brain wide, and on a region by region basis, allowed the identification of biological processes and genes that have diverged in their co-expression relationships over evolutionary time. The preservation of most mouse modules in humans, and many human modules in mouse overall, suggests a basal level of co-expression structure shared between human and mouse, which is consistent with expectations and previous studies (Eidsaa, Stubbs, & Almaas, 2017; Miller et al., 2010; Monaco, van Dam, Casal Novo Ribeiro, Larbi, & de Magalhaes, 2015; Tsaparas, Marino-Ramirez, Bodenreider, Koonin, & Jordan, 2006). However, the strong divergence of several specific human modules from mouse supports acquisition of transcriptomic complexity on the human lineage that is not shared in mouse. Further, module divergence is significantly related to independent measures of selection, including divergence of regulatory and protein coding constraint, indicating that it is capturing evolutionarily relevant characteristics.

The stronger divergence of human modules to mouse than mouse modules to human underlies our observation of “asymmetric divergence” in co-expression relationships. At a cell-type level, this asymmetric divergence is greatest for microglial modules, whereas at a regional level this divergence is greatest in cerebral cortex, an observation consistent with known evolutionary hierarchies (Konopka et al., 2012; Oldham et al., 2006). We use this robustly defined network structure to identify a number of disease associated genes, including PSEN-1, which was previously shown to be divergent based on an independent analysis of microarray data (Miller et al., 2010). Combined with the current analysis, this strongly suggests that mouse models of PSEN-1 will not model human processes with high fidelity; notably mice harboring dominant highly PS-1 mutations do not show frank neurodegeneration or model human AD (Hall & Roberson, 2012). Other disease genes that show lack of conservation of human co-expression relationships in mouse include the ASD risk genes, SCN2A and SHANK3. Divergent genes tended to be expressed at higher levels in human relative to mouse. We observe that this is not broadly due to changes in cell-type composition, but rather generally reflects cellular regulatory effects.

### Assessing evolutionary divergence using co-expression networks

In our analysis, we use module divergence, (a ratio of two species Zsum scores) as a proxy for evolutionary divergence. This metric allows us to overcome the effect of module size on preservation (Fig S1E) and provides a more quantitative basis to compare transcriptome divergence between different processes. Assessing module preservation in non-human primate (NHP) allows prediction of whether transcriptional differences between human and mouse originated before the LCA with NHP, or reflect differences that occurred after the LCA with NHP and are therefore more human specific (Preuss, Caceres, Oldham, & Geschwind, 2004; Varki, Geschwind, & Eichler, 2008). We identify 13 modules where the NHP preservation scores were significantly closer to mouse than human, implicating greater divergence on the human, rather than primate lineage. These modules may therefore contribute to the differentiation between human and NHP brain. In the future, when additional expression datasets are generated from different NHPs, one should be able to create NHP subgroups and further refine when these expression differences were acquired.

Traditional phylogenetic methods utilize mean gene expression derived from all species of interest and use distance-based methods to construct an evolutionary tree (Brawand et al., 2011; Gu et al., 2013). These methods assess the similarity of gene expression between species and construct a phylogeny to minimize the differences in expression according to their distance in the hierarchy. Assessing preservation of co-expression across different species allows us to explicitly assess evolutionary divergence of different biological processes and when on the phylogenetic lineage transcriptomic differences were acquired. Co-expression approaches assess the relationship of each gene to a module eigengene that is generated in a particular species. Therefore, it would be problematic to apply a distance-based approach between NHP and mouse when using values derived from a *human* co-expression network.

By constructing networks in each species and region individually, we can define the biological processes and cell types for each species independently. Other studies have combined expression datasets from different species or regions into a comprehensive expression matrix before constructing a co-expression network (Konopka et al., 2012; Sousa, Zhu, et al., 2017). These studies can highlight species specific processes as co-expression modules are largely driven by genes that display differential expression between species. But, these approaches do not query species differences in co-expression, which may be unrelated to differential expression, and are therefore complementary to the approach taken within our study.

The construction of regional networks yields modules relating to different cell types across regions, which permitted analysis of preservation in co-expression across difference species in cell subtypes. Single-cell sequencing has enabled the detection of numerous cell classes in both human and mouse, permitting identification of species differences at the cell-type level. Whereas cell-type matching between species allows identification of differentially expressed genes between species (Hodge et al., 2019), expression levels themselves may obey a largely neutral model (Khaitovich et al., 2004). Because co-expression reflects functional mechanisms such as co-regulation, changes in network position reflect changes in function (Oldham et al., 2006; Parikshak et al., 2015). In this regard, we observe that human-upregulated genes tend to display stronger kME divergence, consistent with potential adaptive evolution. Still, a substantial portion (73%) of kME divergent genes show similar or reduced expression levels in humans (Fig S4E), more consistent with a model of neutral evolution (Khaitovich et al., 2004). For example, the astrocytic gene, PARD3B, shows stable interspecies expression levels (< 0.5 logFC) in both bulk and single-cell expression data, but shows strong human to mouse divergence at the co-expression level (kME div = 0.51; p < 0.01), which indicates a functional change. On the other hand, IL17D shows considerably higher expression (> 2 logFC) in human bulk and single-cell data, yet is not significantly divergent for co-expression, consistent with a neutral model. Differential expression has been successful in identifying gene expression differences between cell types or brain regions, however preservation of gene co-expression has been suggested to be more successful in recapitulating evolutionary hierarchies (Oldham et al., 2006; Varki et al., 2008) and may therefore be more suited to assessing functional differences between species.

Single-cell sequencing may not detect genes with low levels of expression, which generally reside in the periphery of cell-type co-expression modules (Fig S5A, B). As genes on the periphery of cell-type modules show the greatest divergence of co-expression (Fig S5C-D), until a greater depth can be afforded, co-expression analysis of bulk tissue sequencing will remain important for identifying evolutionary differences, as co-expression analysis of bulk expression data captures genes across a wider range of expression and network position, not just the most central (Masalia, Bewick, & Burke, 2017).

A recent important study used co-expression analysis to identify ‘high fidelity’ markers for broad cell classes across a number of brain regions in both human and mouse (Kelley et al., 2018). Although our identified diverged human-mouse genes are highly overlapping with species specific “high-fidelity” markers (Kelley et al., 2018), our study approached this issue of divergence differently, starting with a discovery dataset, and subsequently assessing co-expression in independent test datasets from different species. Differences in preservation between studies may be due to technical differences, such as RNA-extraction method and sequencing platform, or biological differences such as subject age and housing conditions. Identifying evolutionary differences unrelated to these study differences will therefore increase the signal associated with true evolutionary differences between species. We bootstrapped the effect of study to create confidence intervals around all module and gene divergence scores, allowing us the assess the potential impact of these technical and biological “study” effects.

### Modeling brain function and disease in mouse

Given the ubiquitous nature of mouse in biomedical research for modeling neurological disease (Eppig et al., 2015), it is important to understand species-specific differences. We observe that human glia are highly diverged from mouse suggesting that it may be difficult to make extrapolations of these cell types in human, especially when considering the transcriptome. For example, many transcriptomic perturbations in neuropsychiatric diseases (Fig 5F; (Allen et al., 2016; Gandal et al., 2018)) are associated with immune-glial activation, a response likely to be symptomatic of neuronal dysregulation. Therefore, the initial neurobiological causes of these diseases may be recapitulated in mouse models, but their downstream transcriptional outputs may differ.

By calculating gene-level divergence, we highlight genes that may drive the divergence of cell types and other biological processes in human. For example, ACBD7 and CYBRD1, both in the astrocytic module WB.M6, have both been highlighted in a recent paper to be human-specific astrocyte markers (Kelley et al., 2018), which we confirm in our analysis (kMEdiv >= 0.4; p < 0.01). Kelley et al. (2018) also showed that PMP2, another previously identified human-specific astrocyte gene, when upregulated in mouse astrocytes, was able to increase the number of primary processes and size of mouse astrocytes (Kelley et al., 2018), which is a well-cited distinction between human and mouse (Oberheim et al., 2009). In our dataset, of all genes, PMP2 displayed the greatest change in expression between human and mouse, and showed strong divergence of co-expression in CTX (kMEdiv = 0.54; p<0.01). As supported by single-cell data, PMP2 is associated with both astrocytes (WB.M6; mean kME=0.48) and oligodendrocytes (WB.M7, mean kME=0.29), which may have prevented module assignment in other regions (Hodge et al., 2019). In addition to these specified genes, we identify hundreds of significantly diverged genes for astrocytes and other cell types, for which functional experiments may elucidate the genes effect on making their respective cell type in mouse more “human” (Table S3). For example, as the functional effect of PMP2 upregulation was relatively modest (Kelley et al., 2018), we predict upregulating additional genes may allow human and mouse cell types to become increasingly comparable.

Assessing the top 100 divergent gene pairs from each ‘Whole Brain’ consensus module can highlight novel human-specific functions for each cell type (Table S5). Human-specific associations of glutamate transporters SLC1A3 and SLC1A2 (EAAT1/EAAT2) in the astrocytic WB.M6 module suggest *human* astrocytes have an increased capability to provide glutamate to adjacent neurons. Numerous highly divergent genes of the oligodendrocyte WB.M7 module (e.g. PSEN-1, HSPA2) are associated with Alzheimer’s (AD) (Kelleher & Shen, 2017; Petyuk et al., 2018). Furthermore, strong divergence of WB.M7 gene pairs involved in carnosine (CARNS1, CNDP1) and copper (SLC31A2) metabolism suggest a human-specific role for metal homeostasis in oligodendrocytes. Given that metal dyshomeostasis induces many of the cognitive symptoms observed in AD, the human-specific nature of metal homeostasis in oligodendrocytes may explain why many treatments derived in mouse have failed in human clinical trials (Ayton, Lei, & Bush, 2015). Additionally, the AD risk gene TREM2 (Pottier et al., 2013) resides amongst the highly divergent microglial WB.M10 genes. The complement cascade genes C1QA-C, C3 and C3AR1 also form many of the highly divergent gene pairs within the microglial WB.M10 module, suggesting a human-specific role for complement mediated synaptic pruning in microglia which may have implications in AD pathology (Hansen, Hanson, & Sheng, 2018; Hemonnot, Hua, Ulmann, & Hirbec, 2019).

We identify dozens of genes currently associated with risk for neurodegenerative and neurodevelopmental disorders whose co-expression is significantly diverged from mouse (Table S7). Remarkably, alpha-synuclein (SNCA), a PD risk gene, shows divergence primarily in the substantia nigra – the first region to display degeneration in PD patients (Surmeier, Obeso, & Halliday, 2017). Presenilin-1 (PSEN-1), an AD risk gene, displayed divergence to mouse across numerous brain regions, but was preserved in NHP. PAX6 and ERLIN2, in the astrocyte module WB.M6, are both implicated in intellectual disability and display co-expression divergence across all cerebral cortex regions. SHANK3 was among 57 other ASD risk genes (Methods; (Ruzzo et al., 2019; Sanders et al., 2015; Satterstrom et al., 2020), displaying significant kME divergence in at least one brain region and we provide a full list in Table S7. Notably, when we broaden the analysis to include the co-expression module representing synaptic vesicle trafficking in which SHANK3 is a core member (BRNHIP.M5; kME=0.91; 28^th^ percentile), we find that 27 out of the 200 genes in that module, including SRF, DOC2B, and LRP3 (Table S9) are also significantly divergent between human and mouse, providing further evidence that some of the basic biological pathways in which SHANK3 participates are also divergent. Together, these findings demonstrate a number of genes that contribute to human disease, but whose function is not likely to be faithfully recapitulated in mouse.

### *In vitro* models of human brain and cell types

Recent advances in *in vitro* modeling of human brain offers the potential to model human brain function in a dish (Kanton et al., 2019; A. M. Pasca et al., 2015; Sloan et al., 2017; Yoon et al., 2019). Cortical organoids faithfully recapitulated astrocytic, activated glial and most neuronal *in vivo* co-expression signatures. Although oligodendrocyte and homeostatic microglial signatures were not captured, future modules should attempt to incorporate these cell types appropriately (Marton et al., 2019; Ormel et al., 2018). Currently, given that aging mouse brain most successfully recapitulated the microglial co-expression signature, some microglial related processes may be more appropriate to study in aging mice, but once microglia are faithfully incorporated into organoid models, their preservation should be carefully tested (Abud et al., 2017; Ormel et al., 2018). The most notable advantage of cortical organoids compared to mouse is the faithful recapitulation of human astrocytes, which appear to model human astrocytes similarly to NHP *in vivo*. For example, ARHGEF6, a member of the astrocytic WB.M6 module is associated with X-linked mental retardation and is significantly more preserved in organoids than mouse, making organoids a preferred model to study mechanisms underlying this disease.

Interestingly, astrocyte (and oligodendrocyte) markers derived from sorting experiments (Zhang et al., 2016) did not show strong divergence in co-expression from human to mouse (Fig S2D). These cell types were sorted based upon HepaCam and GalC markers respectively and therefore may not have captured all glial sub-populations, some of which perhaps representing more highly diverged, human specific aspects of glial biology. Alternatively, immunopanning and culturing of human astrocytes may remove them from their physiologically optimal state in a 3D environment and cause them to lose their human-specific properties, both transcriptionally and functionally. Interestingly, mice with brain-engrafted human glial progenitors and astrocytes displayed enhancement of both activity-dependent plasticity and learning (Han et al., 2013). So, although physiological environment may be important for astrocytes to manifest their human-specific components, this environment may be somewhat shared between human brain, cortical organoids and mouse brain.

### Limitations and further work

This study highlights a number of transcriptomic differences between species, especially for glial cell types. Most identified differences are likely due to evolutionary differences between species, however, we cannot exclude the effect of external confounding factors such as environment, diet or agonal state. For example, given the sterile housing conditions of mice, we hypothesize that immunological differences in human could be due to non-sterile conditions. We cannot rule out some contribution of environmental differences to the divergence of this activated glial signature. But it is important to note that regardless of cause, this cell state is not captured in mouse. Mitigating against a major or pervasive contribution of environmental effects to these differences, we find that co-expression divergence was strongly correlated with sequence divergence, which would drive the differential regulation of gene expression (Khaitovich et al., 2005; Monaco et al., 2015). Moreover, we observed that this activated microglial signature was not specific to humans, but also was observed in NHP housed in sterile conditions.

To assess module preservation, we utilize the combination of many test expression datasets that were not assayed uniformly across all brain regions in this study. Therefore, module preservation of each brain region may utilize a different combination of test datasets, which may differ by sample preparation, developmental time point, or environmental state. To assess the effect of study selection upon regional divergence, we regressed out any “study-specific” effects upon module divergence and observe that regional divergence after study regression is highly correlated the raw regional divergence scores (Fig S1F). This suggests the non-uniform distribution of test expression datasets across brain regions does not bias regional preservation scores, although there may still be a small confound between brain region and factors underlying study design. We perform study-level permutations to calculate region-specific divergence differences to further account for the variability in study choice to mitigate this issue.

This study provides a multi-region, multi-species comparison of the evolutionary divergence of transcriptomic networks generated from adult human brain. However, the brain also exists under a number of different developmental states or environmental conditions, which would need to be further investigated to achieve a more complete understanding of species differences. However, these analyses, based on dozens of data sets, and multiple brain regions, provides a robust framework for understanding major species differences.

## Methods and Materials

### Acquisition of expression datasets

To assess the evolutionary conservation of brain networks, we processed 7,287 samples from 12 brain regions in human – cerebellum (CBL), cerebellar hemisphere (CBH), dorso-lateral pre-frontal cortex (CTX), Brodmann area 9 (CTXBA9), Brodmann area 24 (CTXB24), hippocampus (HIP), amygdala (AMY), hypothalamus (HYP), substantia nigra (SNA), nucleus accumbens (ACC), caudate nucleus (CDT), and putamen (PUT), 2,933 samples from six brain regions – cerebellum (CBL), cortex (CTX), hippocampus (HIP), amygdala (AMY), hypothalamus (HYP), and basal ganglia (BG), of three non-human primates (NHP) - macaque, baboon and chimpanzee and 6,667 samples from seven brain regions - cerebellum (CBL), cortex (CTX), hippocampus (HIP), amygdala (AMY), hypothalamus (HYP), nucleus accumbens (NACC), and striatum (STR), in mouse (Table S1). We also measured network preservation against *in vitro* brain organoid systems from eight independent studies (Table S2). Only organoids older than day 50 were used for this analysis as a compromise between statistical power and accurate matching to mature human brain. Individual studies are listed in Supplementary Table 2. Given the lack of independent hypothalamic expression datasets in NHP, this brain region was omitted from NHP-related analysis.

### Processing of gene expression data

Unless specified, normalized expression values were obtained from GEO or from the study authors directly. For mouse RNA-seq data, samples were aligned to the GrCm38 transcriptome using Salmon producing TPM values. Genes were retained if they had >20% non-zero values for each study of a brain region and were subsequently log2 (+0.001) transformed. To ensure co-expression was not driven by outlying samples, samples were removed if they had a low connectivity to the remaining samples (z.k < -2.5 IAC), or if they displayed over 4 SDs from the mean for any of the top 10 expression PCs. Outlier removal was iterated up to 5 times or until <3 samples were removed in an iteration. All available technical (ie. batch, alignment statistics) and biological covariates (ie. age, sex, sub-region) from the expression data were included in the linear regression providing that the covariate explained on average over 1% of the expression variance, while ensuring that at least 8 degrees of freedom remained in the dataset. For samples quantified in-house, both STAR and picard covariates were log2 (+ 0.001) transformed and combined to make a set of sequencing PCs that were included as possible technical covariates for regression. If the expression dataset sample size was <10, covariates were not regressed. To allow comparison between human and mouse datasets, gene IDs were converted to their corresponding human/mouse orthologue Ensembl gene ID using the biomaRt package in R.

### Network generation

To generate human and mouse brain networks, we utilized robust WGCNA on 12 brain regions in human using RNA-seq data from the GTEx consortium (Consortium et al., 2017) and 7 brain regions in mouse using a number of publically available RNA-seq datasets (Table S2). To prevent the effect of study driving mouse network construction, we removed batch effects between study using ComBat (Leek, Johnson, Parker, Jaffe, & Storey, 2012) before combining datasets into a single expression matrix. Each regional network was constructed using the consensus of 50 bootstrapped networks. Regional networks were then merged in a hierarchical manner until a Whole-Brain (WB) consensus network was generated (Hartl et al., 2020). In mouse, the Whole-Brain consensus network was created using a consensus of the 7 region-specific networks, having subset for common genes.

To generate co-expression modules in mouse, we selected a deepsplit cut height that gave the number of modules closest to 20. We removed modules which had limited support from independent microarray datasets (Zsum UQ <5). For each region, the nomenclature of each WB module would be adopted by the regional module that shared the greatest similarity (Jaccards Index > 0.4). The remaining regional modules were numerically named according to module size, for instance AMY.1 and AMY.6 being the largest and smallest amygdala modules respectively that did not show significant similarity to a WB consensus module.

### Module annotation

For cell-type enrichment, a hypergeometric test was performed for each module against cell-type marker gene lists derived from sorting experiments (Cahoy et al., 2008; Holtman et al., 2015; Mancarci et al., 2017; Zhang et al., 2014; Zhang et al., 2016). A module was assigned to the cell-type for which it had the strongest enrichment, providing the odds ratio (OR) was above 3 and did not show stronger enrichment for mitochondrial or ribosomal related genes. Gene ontology was assessed using the R package gProfileR (Reimand, Kull, Peterson, Hansen, & Vilo, 2007).

### Calculating module preservation scores

To query the extent to which a module was reproducible (preserved), every module of each reference network was tested against many independent expression datasets derived from the same, or highly similar, brain region for each species (Table S1). Preservation analysis was also performed using marker gene lists in addition to co-expression modules (Table S10). For analysis comparing young and aging adult brain, samples were partitioned into two groups (young and aging adult) for each study (n=6 human; n=5 mouse) on a regional basis. Young adult is defined as 13-40 years old in human and 2-14 months old in mouse. Aging adult is defined as 40+ years in human and 14+ months in mouse. Module preservation was executed using 25 permutations. For each test, a composite Zsummary (Zsum) statistic was generated. The larger the z-score the more likely it is to be preserved in the test dataset, traditionally a Zsum score >10 indicates strong module preservation whereas a score <2 suggests weak preservation. Zsum scores were capped at 40 to prevent inflated aggregate preservation scores. The Zsummary score is generated from a number of other statistics such as module density (how densely connected the module genes are in the test network) or connectivity (how well connected the module genes are in the test network). For further details, see (Langfelder et al., 2011). The Zsum score shows a strong dependence on module size – with larger modules having greater statistical significance to be called preserved than smaller modules (Fig S1E). However, this dependence on module size does not affect module divergence scores as these are calculated as a ratio of two preservation scores.

### Calculating module divergence scores

The upper quartile (UQ) of all Zsum preservation scores was calculated for human, non-human primate (NHP) and mouse independently, generating UQ hZsum, UQ pZsum and UQ mZsum scores respectively. The UQ was chosen to compare between evolutionary classes in order to remove emphasis from lowly preserved datasets, perhaps collected from an unrelated developmental stage or differing expression profiling platform. Modules with a same-species UQ Zsum score of over 5 were deemed reproducible and retained for further analysis (Fig 1B). Comparing the UQ Zsum scores between species allows us to determine module-level co-expression differences between species. To assess divergence for each human and mouse module, the fold change (FC) difference was calculated between the UQ hZsum and UQ mZsum scores (Fig 2A). These divergence scores were capped at a score of 15 to remove the effect of extreme outliers.

For all human modules, to gain a greater understanding of when these transcriptomic changes were acquired, we were also able to leverage the UQ pZsum scores to create a 1) ‘human-specificity’ (HS) score, where transcriptional changes occurred on the human lineage after divergence with the last common ancestor (LCA) of NHP, and a 2) ‘primate specificity’ (PS) score, where changes occurred before divergence with the LCA of NHP (Fig 4A). Human specificity can also be interpreted as Human-Primate divergence, whereas primate specificity interpreted as Primate (including human) - Mouse divergence. The HS score is the fold-change difference between the UQ hZsum score and the UQ mZsum or UQ pZsum score (whichever was larger). The PS score is the fold-change difference between the UQ hZsum score or UQ pZsum score (whichever is smaller) and the UQ mZsum score. These calculations can be summarized as below.

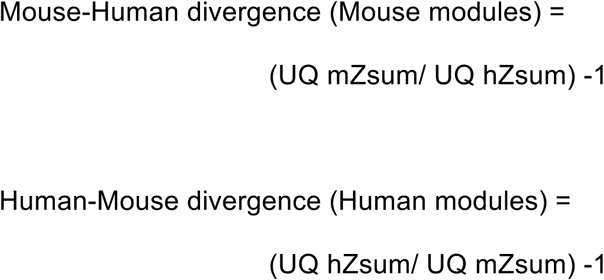

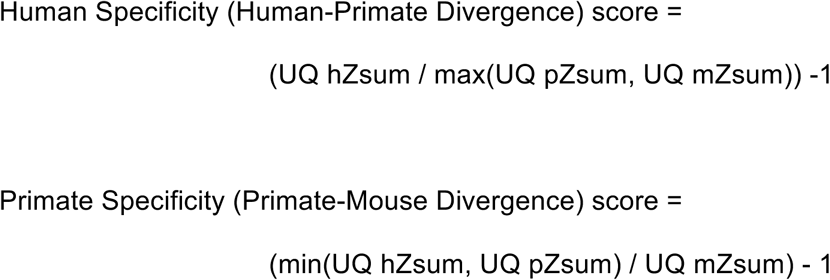

### Calculating gene-level divergence scores

To identify gene drivers of module divergence we calculated the correlation of each gene to its module eigengene (kME) in the discovery dataset and all test datasets in human, NHP and mouse. Next, we calculated the kME difference between each species and the discovery dataset. The difference in mean kME values between human and mouse was used to calculate a kME divergence score, which was used to highlight highly divergent genes between species. For each region, kME values and divergence scores were generated for both regional and Whole-Brain (WB) modules

### Calculating human mouse expression differences

To assess the difference in mean expression between human and mouse, both bulk and single-cell data were used. For bulk comparison, TPM values for each gene using the human (GTEx) and mouse (compiled) discovery RNA-seq expression data on a region by region basis. The human mouse expression fold-change difference was calculated as (Human TPM + 0.01) / (Mouse TPM + 0.01). To calculate a consensus expression difference between human and mouse, TPM values were averaged across regions before calculating fold-change.

To assess the change in expression between human and mouse, while matching for cell type, we utilized single-cell expression data derived from human and mouse cortex (Hodge et al., 2019). For human and mouse respectively, we utilized the available trimmed-mean and median expression TPM values for all genes in each of their identified cell-type clusters. For human, 120 cell-type clusters were identified, for which we assigned 111 clusters into a neuronal group, 3 clusters into an astrocyte group, 3 clusters into an oligodendrocyte group, 1 cluster into a microglial group and 1 cluster into an endothelial group. For mouse, 290 cell-type clusters were identified, for which we assigned 271 clusters into a neuronal group, 2 clusters into an astrocyte group, 6 clusters into an oligodendrocyte group and 3 clusters into an endothelial group. To compare the human mouse expression fold-change difference for each cell type, we calculated the mean expression across all clusters within a cell-type group and compared the mean expression between species. To compare the human mouse FC difference between bulk and single-cell data, genes with co-expression membership to cell-type modules were compared with their respective cell-type in single-cell expression data.

### Disease enrichment

To assess enrichment of disease relevant genes in human-mouse diverged genes, a hypergeometric test was performed for diverged genes against the top 500 genes up and down regulated in autism, schizophrenia, bipolar disorder, depression, anxiety and Alzheimer’s (Allen et al., 2016; Gandal et al., 2018).

To investigate genes with causal association to human disease we curated: a) genes harboring high risk likely protein-disrupting mutations in ASD patients (Ruzzo et al., 2019; Sanders et al., 2015; Satterstrom et al., 2020), b) genes of SFARI categories 1 or 2 (Abrahams et al., 2013), c) genes within a nervous system disease KEGG pathway (Kanehisa & Sato, 2020), d) genes of the DisGeNet database score linked to mental or behavioral dysfunction with an association score above 0.5 (Pinero et al., 2017; Pinero et al., 2020) and e) previously curated genes implicated in neurodegenerative diseases (Fu, Hardy, & Duff, 2018).

### Calculating sequence divergence

To measure sequence divergence of genes in each module the 3’ UTR, 5’ UTR, promoter (250 / 2000 bp upstream of transcription start site), introns, exons and splice sites of each gene was determined and PhastCons score generated. The PhastCons score is a measure of DNA sequence conservation across 40 different mammalian species with constrained sequences obtaining a higher score (Siepel et al., 2005). The median gene PhastCons score was calculated for each module – providing a module level sequence divergence score for many gene annotations.

### Availability of Data and Materials

The datasets supporting the conclusions of this article can be accessed through the GEO or other online repositories with unique identifiers listed in Table S2.

## Acknowledgements

We are grateful for feedback and discussion from GR, JR and members of the Geschwind lab. This research was supported by the NIMH grant 5R01MH109912 and NIH grant 5U01MH115746.

## Author contributions

WGP and DHG designed the experiments and approach, and edited the manuscript. WGP wrote the first draft of the manuscript, performed data analysis and developed the methodological approaches under supervision of DHG. CHH aided in data analysis and analytical strategies.

---

**Figure S1.**
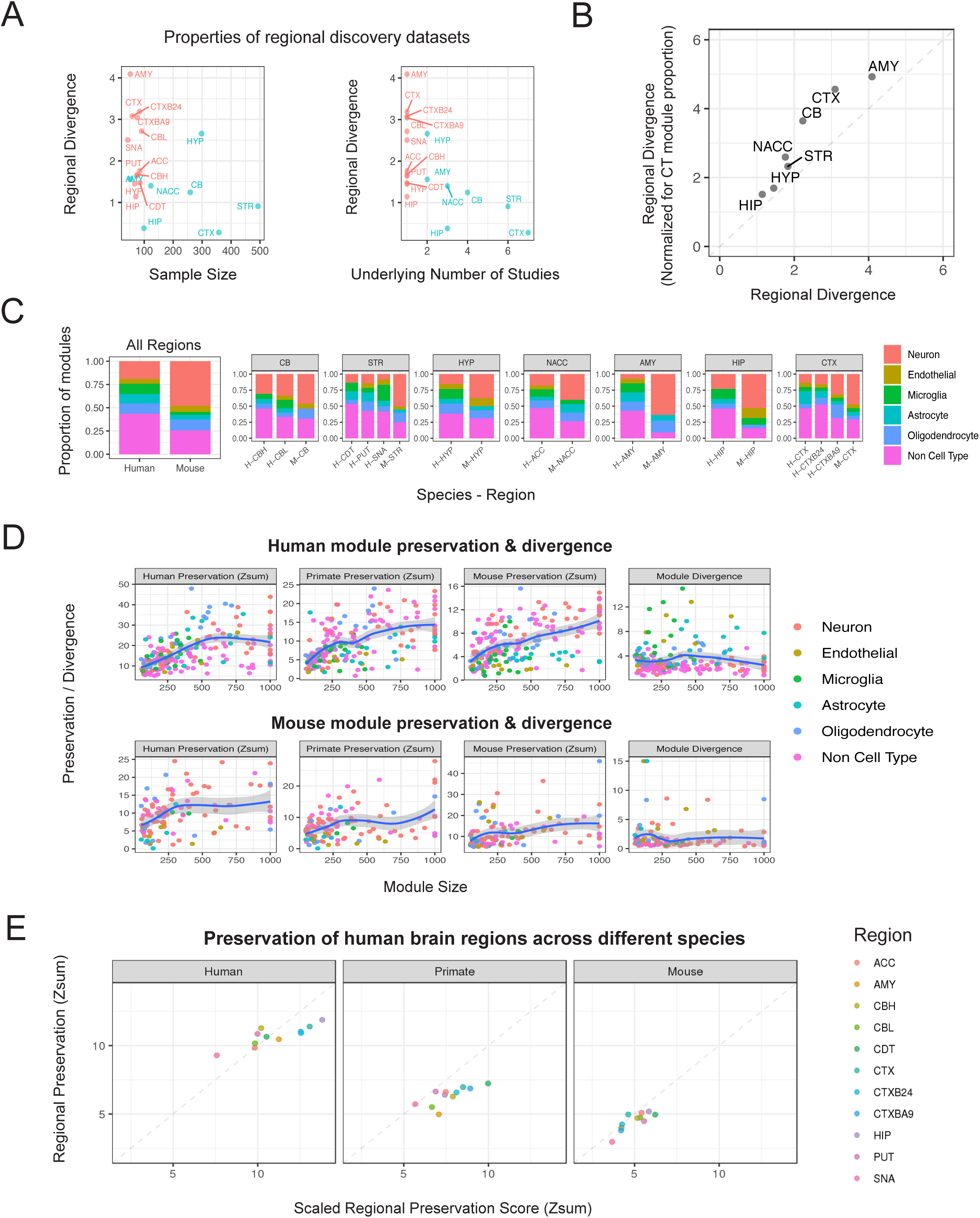

**Figure S2.**
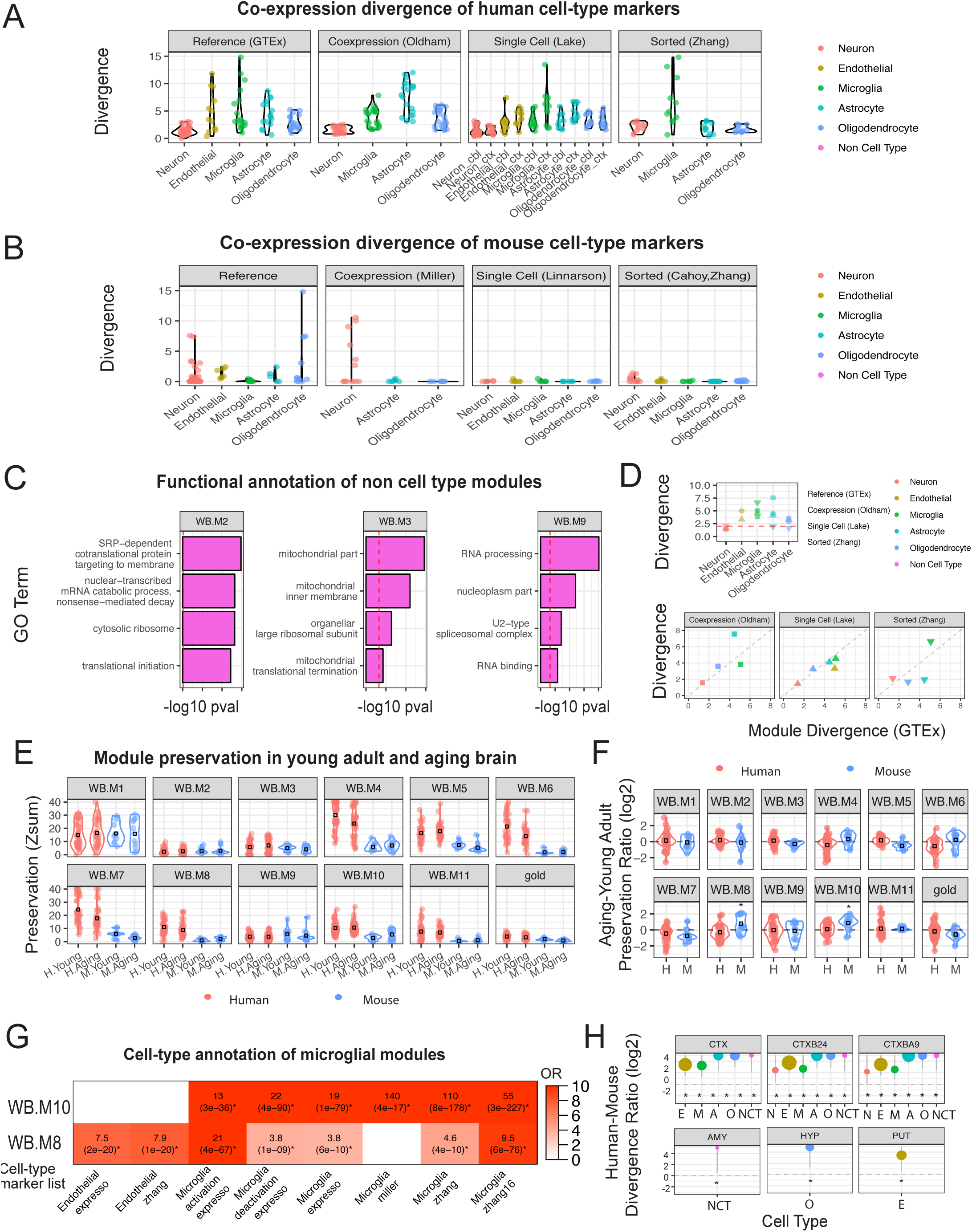

**Figure S3.**
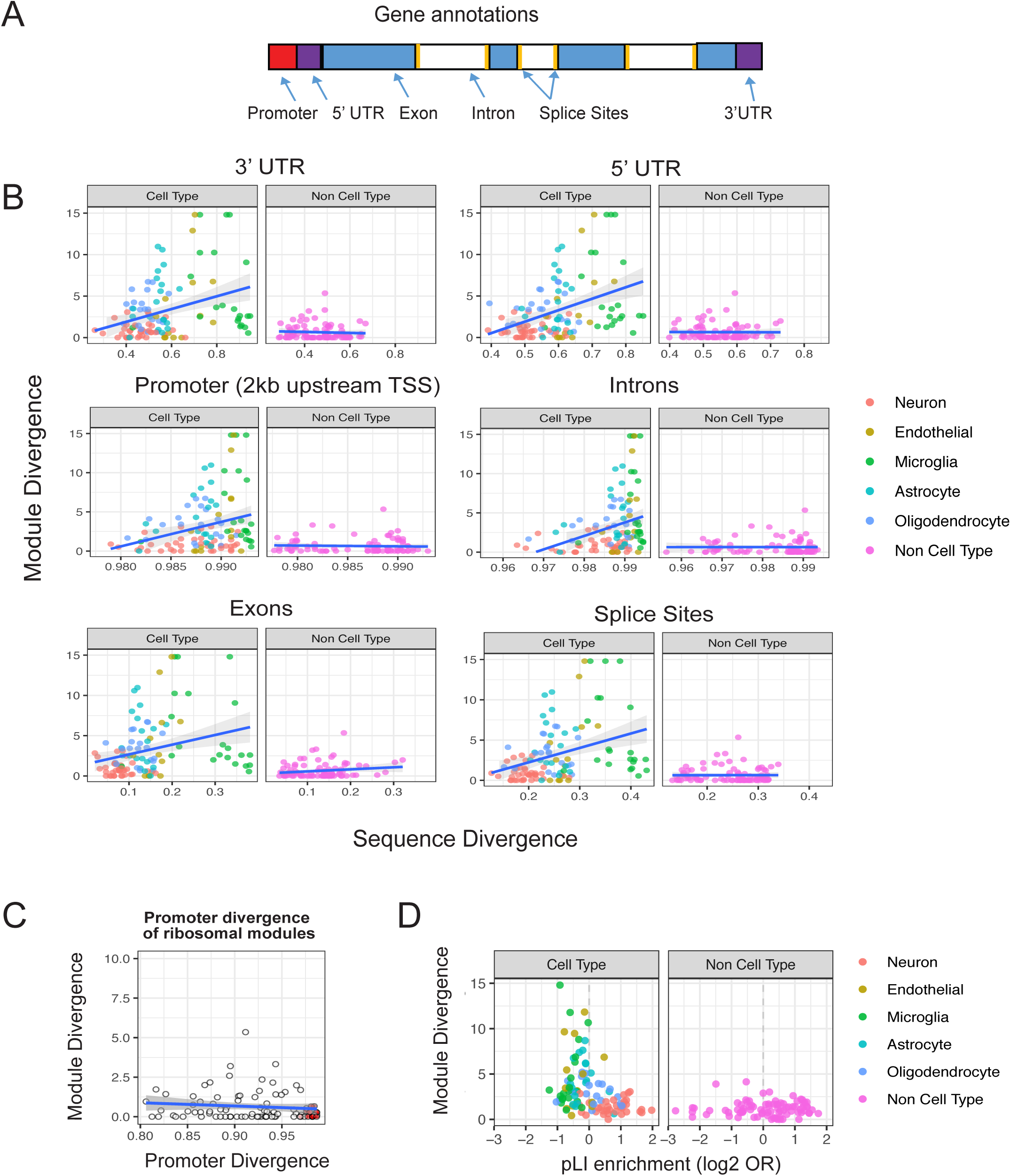

**Figure S4.**
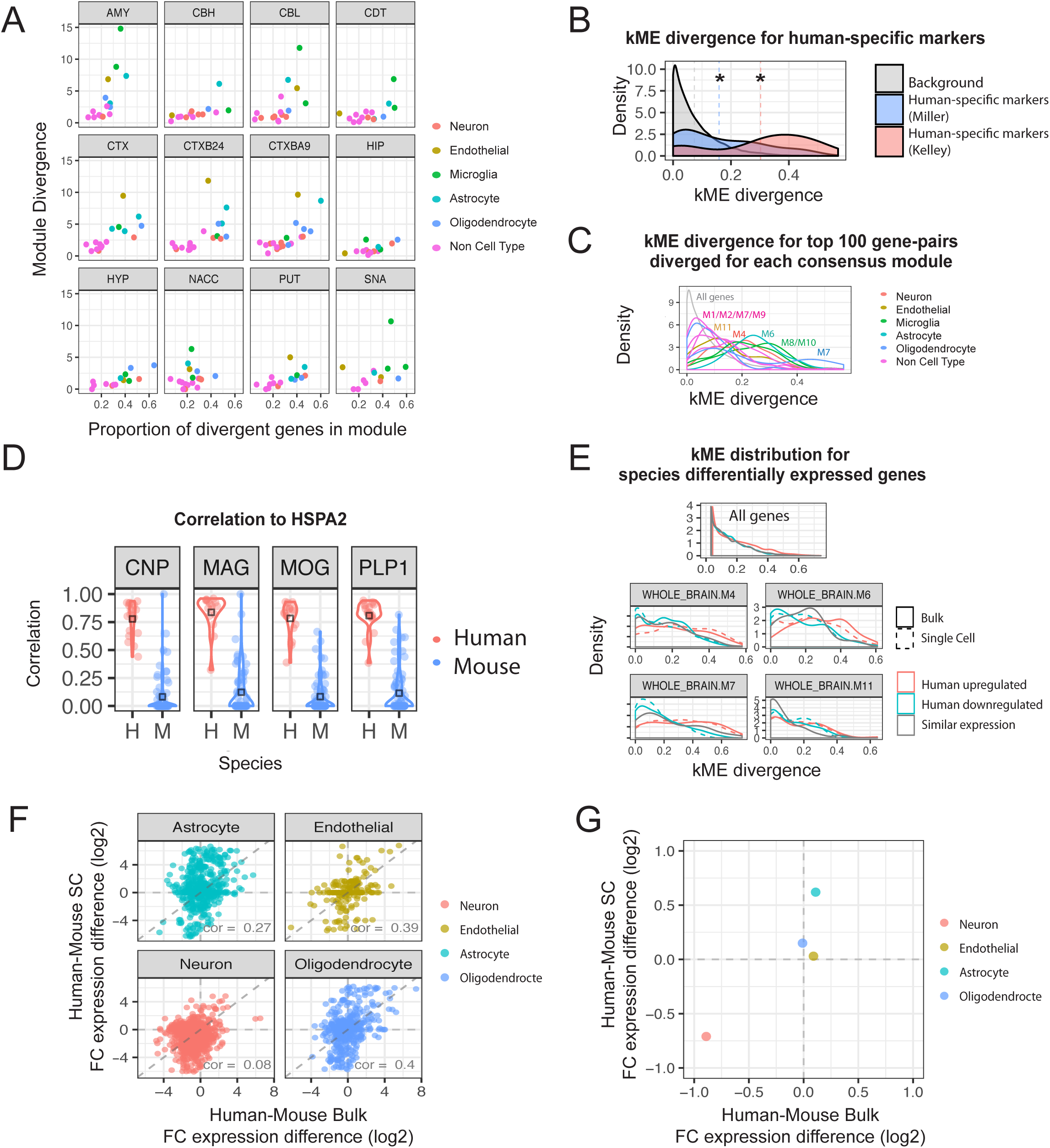

**Figure S5.**
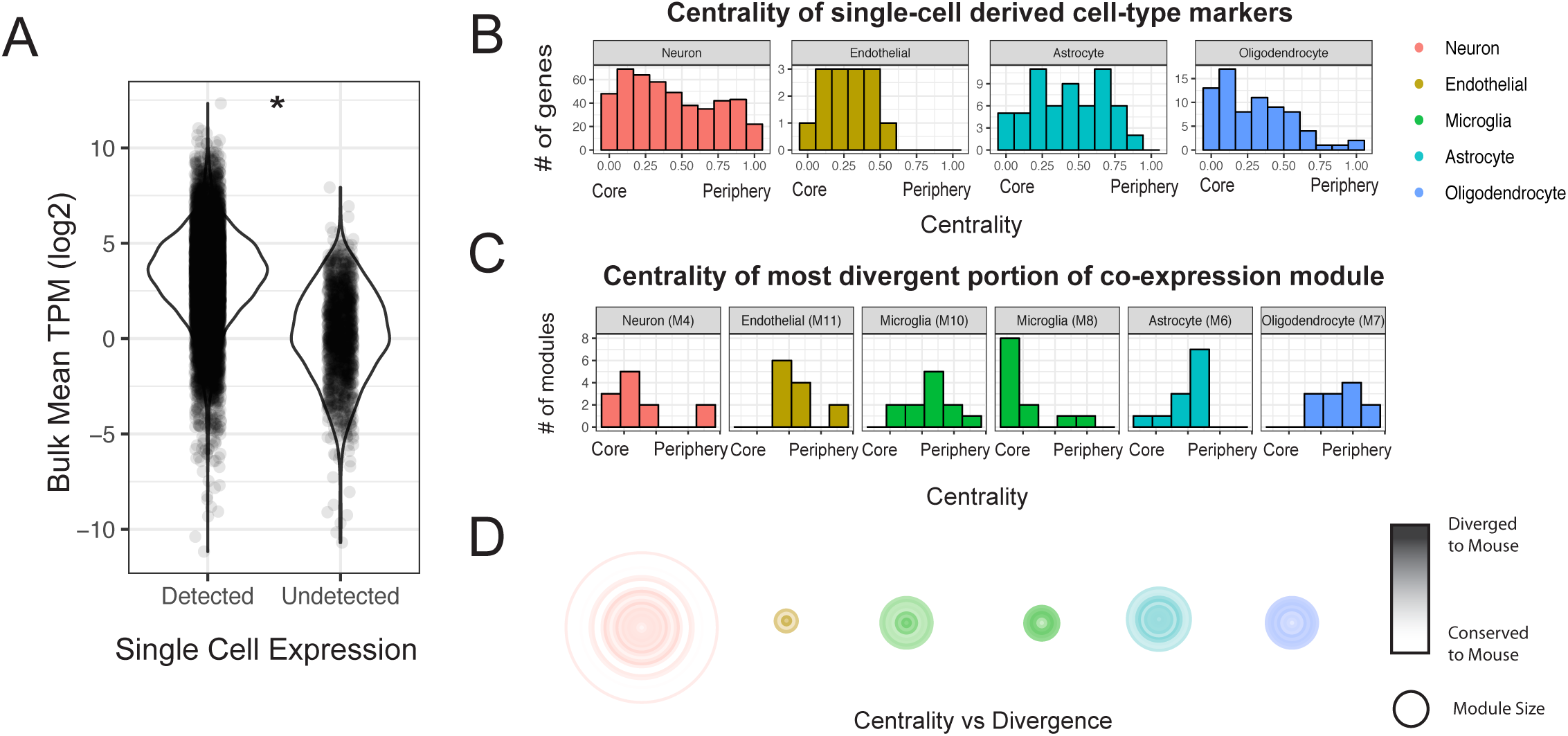

